# Acoustic Characteristics of Low-Frequency Vocalizations of Bryde’s Whales (*Balaenoptera edeni edeni*) in the Weizhou Island Waters of the Beibu Gulf, China

**DOI:** 10.64898/2026.06.17.729802

**Authors:** Yining Yang, Xingyu Li, Make Li, Mo Chen, Fei Fan, Kexiong Wang, Hao Du

## Abstract

Bryde’s whales (*Balaenoptera edeni edeni*) are a first-class nationally protected species in China, and the waters around Weizhou Island in the Beibu Gulf support one of the country’s few regularly observed coastal groups. Acoustic data for this population are sparse, and vessel noise here has not been measured. We conducted 16 vessel-based surveys around Weizhou Island between 11 and 31 January 2024 from a low-disturbance sailboat platform, with concurrent visual observation, and measured 729 low-frequency signals attributed to Bryde’s whales. All signals were pulsed, with a repetition rate of 20–56 pulses s⁻¹, and their energy (400–1100 Hz) places them among the highest-frequency Bryde’s whale calls reported. Signal duration was negatively correlated with maximum and center frequency in linear mixed-effects models that accounted for the non-independence of signals recorded within a session, but was unrelated to minimum frequency or bandwidth. Clustering of six standardized parameters, supported by bootstrap resampling (mean adjusted Rand index 0.87) and a Gaussian mixture model (minimum BIC at three components), resolved three call types differing in onset frequency, bandwidth and duration that co-occurred across sessions. In structure and frequency range these calls most closely resemble the pulsed moans and discrete pulse series described from juvenile Bryde’s whales and mother–calf pairs, and they are distinct from the lower-frequency tonal calls (25–270 Hz) reported from the same region. Speedboat passage raised levels by 12.5 dB over the 20 Hz–20 kHz band and by 8.0 dB within the 400–1100 Hz band, cutting the detection margin of the calls from 10 dB to about 2 dB. The implied source levels match small high-speed craft elsewhere, so the masking risk is regional rather than local.

## 1 Introduction

Bryde’s whales (*Balaenoptera edeni edeni*) are listed as first-class protected wildlife in China. Weizhou Island in the Beibu Gulf is one of the few places where a coastal group is seen regularly, and whale-watching speedboats now run trips around these animals on most days. Published acoustic data for the population are sparse, and the noise those boats add to the habitat has not been measured.

Recent studies have described the spatiotemporal distribution, habitat use and foraging behaviour of Bryde’s whales around Weizhou Island by integrating citizen-science reports, questionnaire surveys, ship-based observations, satellite remote sensing, molecular identification and photo-identification (Chen et al., 2019; Zhang et al., 2021; Liu et al., 2021; He, 2021; Wu, 2021; Sun et al., 2024). Few of them recorded underwater sound. Previous work has shown that vessel noise can alter baleen whale behaviour and vocal output, including through noise-related changes in call amplitude consistent with the Lombard effect (Blair et al., 2016; Helble et al., 2020; Johnston and Painter, 2024; McKenna, 2011; Rolland et al., 2012). For this population, the only published acoustic measurements are the low-frequency tonal calls reported by Wang et al. (2022) at 39–268 Hz. Here we used a sailboat-based survey platform and passive acoustic recording to (i) characterize the spectral and temporal parameters of low-frequency Bryde’s whale vocalizations, (ii) test whether these signals form distinguishable call types, (iii) describe within-session calling rates, and (iv) quantify the change in underwater sound levels associated with speedboat passage and the spectral overlap with the recorded calls.

## 2 Materials and Methods

### 2.1 Study period and location

The study was conducted around Weizhou Island and adjacent waters in the central Beibu Gulf, Guangxi, China. Weizhou Island lies northwest of Xieyang Island, and offshore oil and gas platforms are located to the southwest. Surveys were conducted from 11 to 31 January 2024 (21 days, 16 survey days). The survey area included waters within 12 nautical miles of Weizhou Island, waters around Xieyang Island, the channel between Weizhou and Xieyang Islands, and waters adjacent to the Southwest Oil and Gas Field.

### 2.2 Field survey and acoustic recording

Surveys were conducted from a sailboat platform (Figure 1a). Acoustic recordings were made with a SoundTrap 300HF recorder (Ocean Instruments Ltd., New Zealand), a self-contained passive acoustic monitoring instrument with an integrated hydrophone, preamplifier, recording unit, memory and battery. The recorder has a frequency response of 20 Hz to 150 kHz, a sensitivity of −203 dB re 1 V/µPa, two gain settings, a maximum sampling rate of 576 kHz and 16-bit analogue-to-digital conversion. Recordings used in the present analysis were acquired at a sampling rate of 576 kHz with the high-gain setting and the high-pass filter disabled, as documented in the instrument log files. A DolphinEar PRO real-time hydrophone was used for auxiliary monitoring. This hydrophone is rated to 100 m depth, operates from 1 to 24,000 Hz and connects through a three-pin XLR interface. Equipment was configured for continuous recording to maximize the probability of detecting whale vocalizations for subsequent analysis.

**Figure 1.**
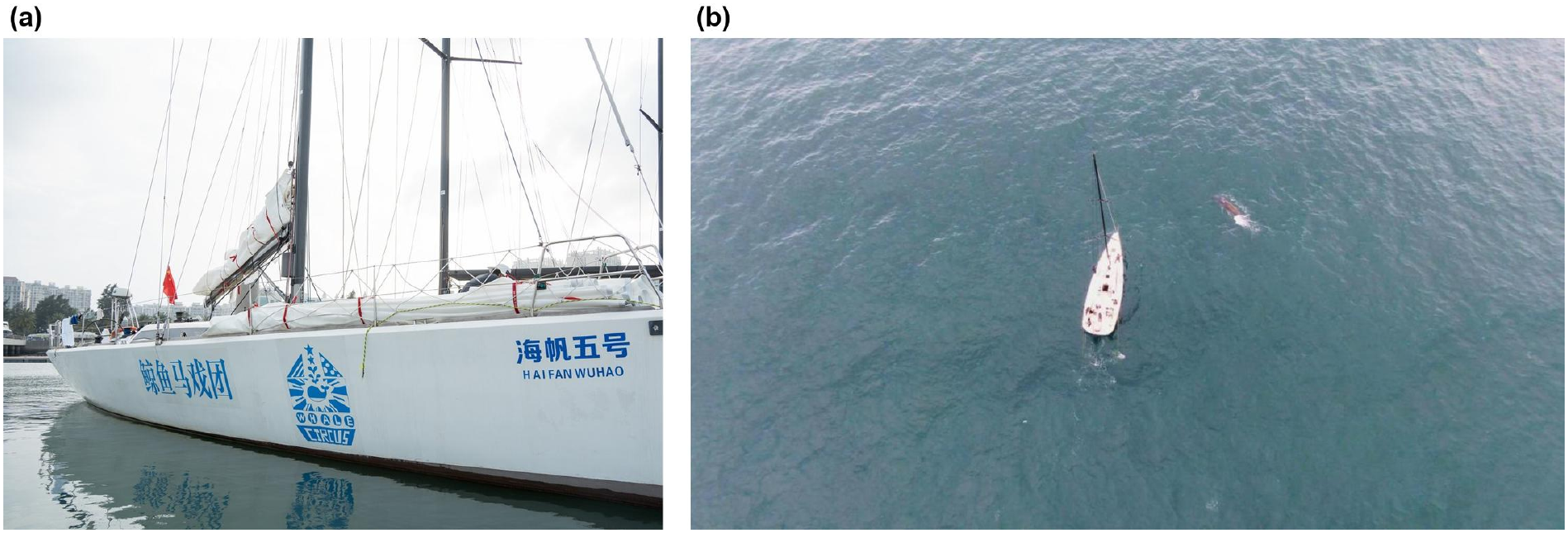
The survey platform. (a) Exterior view of the sailboat used for all surveys; (b) aerial view of the vessel with a Bryde’s whale alongside.

Signals recorded on 21 January were acquired with a second, independent recording system operating at a sampling rate of 96 kHz (one session mono/16-bit; one session stereo/24-bit) as part of a parallel survey effort. These recordings used a different device, hydrophone and calibration chain, and are therefore identified as a distinct recording configuration in all analyses (Table 1); the implications of pooling them with the SoundTrap data are addressed in the Discussion.

**Table 1.** Recording sessions and detection summary.

| Recording session | Date | Recorder | Signals (n) | Analysed span (min) | Call rate (min<sup>−1</sup>) | Evidence category |
|---|---|---|---|---|---|---|
| S1 | 13 Jan (AM) | SoundTrap 300HF, 576 kHz | 46 | 32.8 | 1.40 | acoustic-only |
| S2 | 13 Jan (PM) | SoundTrap 300HF, 576 kHz | 16 | 3.3 | 4.92 | video-confirmed |
| S3 | 14 Jan (night) | SoundTrap 300HF, 576 kHz | 1 | — | — | acoustic-only |
| S4 | 15 Jan (AM) | SoundTrap 300HF, 576 kHz | 40 | 7.1 | 5.60 | video-confirmed |
| S5 | 16 Jan (PM) | SoundTrap 300HF, 576 kHz | 4 | 17.2 | 0.23 | video-confirmed |
| S6 | 17 Jan (day) | SoundTrap 300HF, 576 kHz | 115 | 304.9 | 0.38 | acoustic-only |
| S7 | 19 Jan (PM) | SoundTrap 300HF, 576 kHz | 334 | 149.0 | 2.24 | visual-only |
| S8 | 20 Jan | SoundTrap 300HF, 576 kHz | 116 | 25.1 | 4.62 | video-confirmed |
| S9 | 21 Jan (AM) | second system, 96 kHz | 9 | 81.4 | 0.11 | visual-only |
| S10 | 21 Jan (PM) | second system, 96 kHz | 48 | 47.4 | 1.01 | acoustic-only |
| **Total** |  |  | **729** |  |  |  |

For deployment, two high-strength nylon ropes, each approximately 20 m long, were prepared with diving weights attached to one end as sinkers. The SoundTrap 300HF was fixed to the rope approximately 1.5 m above the sinker, and the other end of the rope was secured to the stern rail. Hydrophones were deployed only when sea state and vessel movement allowed suitable recordings. When the vessel was moving rapidly under sail, hydrophones were kept out of the water because flow noise and impact with the water column could mask animal signals. When vessel speed was below 2 knots, the recorder was deployed with the rope fully extended. After the sails were furled and the vessel drifted, the rope was shortened to maintain the hydrophone at approximately 2–3 m depth. When the engine was used in calm conditions or during rapid manoeuvring, the hydrophone was retrieved before operation to prevent equipment damage and reduce recording interference. At offshore anchorages in areas frequented by Bryde’s whales, the hydrophone was also deployed at night to record potential vocal activity.

The sailboat was capable of overnight operation at sea, and the captain determined whether daily survey conditions were safe. The vessel departed Nanwan Port, Weizhou Island, at dawn and surveyed until nightfall, when visual detection of Bryde’s whales was no longer reliable. The captain and seven trained crew members conducted visual observations, photographic documentation, sail handling, and hydrophone deployment and retrieval. One crew member served as the primary observer, while other crew members conducted visual observations when not assigned to vessel or equipment duties.

When Bryde’s whales were sighted near the sailboat, the vessel remained stationary and the pre-activated hydrophone was deployed as described above. At the same time, two crew members deployed the real-time hydrophone to monitor for possible whale vocalizations. If whales were detected farther from the vessel, the captain determined whether the sailboat could approach safely under sail. Engine use was avoided where possible to minimize disturbance and reduce interference in the acoustic recordings. Upon visual confirmation of whale location, time and position were recorded for later comparison with acoustic data.

### 2.3 Acoustic analysis

Compressed audio files (SUB format) recorded by the SoundTrap 300HF were transferred to a workstation and decompressed to WAV format using SoundTrap Host software. WAV files were imported into Raven Pro 1.65 (Cornell Laboratory of Ornithology, Ithaca, NY, USA) for analysis. Spectrograms were displayed using a 65,536-sample Hanning window, corresponding to a frequency resolution of 8.79 Hz, with 90% overlap (time grid 11.4 ms) at the 576 kHz sampling rate. Background environmental recordings were reviewed to assess potential mechanical or other noise interference.

Low-frequency signals attributed to Bryde’s whales were annotated without prior reference to whale sighting timestamps; each audio segment was examined independently, and putative Bryde’s whale signals were tallied. The marine environment included water movement and other background sounds, so subsequent measurements focused on low-frequency signals with a signal-to-noise ratio greater than 10 dB. During annotation two spectrogram views were displayed simultaneously, and selections were therefore exported to two parallel selection tables; duplicate records sharing identical begin/end times and frequency bounds were removed before analysis. One degenerate point-selection with zero duration and zero bandwidth was also excluded. The final annotated dataset comprised 712 signals.

For each signal we measured the following parameters in Raven Pro: duration (Delta Time, the time between selection onset and offset); minimum frequency (MinF, the lower frequency bound of the selection); maximum frequency (MaxF, the upper frequency bound); center frequency (the frequency that divides the selection’s energy into two equal halves); 50% bandwidth (BW50) and 90% bandwidth (BW90), the bandwidths containing 50% and 90% of the selection’s energy, respectively; and delta frequency (MaxF − MinF). Delta frequency is by definition a linear combination of MinF and MaxF and is therefore statistically redundant; it is reported here only for comparability with previously published datasets.

A further 20 signals had been clipped and archived during the field campaign but were never annotated. These were measured with an automated pipeline that reproduced the annotation workflow: mean power spectra were computed with a 65,536-point Hanning window and 90% overlap, the frequency bounds were placed 12 dB (lower bound) and 10 dB (upper bound) below the peak of the baseline-subtracted spectrum, and the center frequency and bandwidths were obtained as energy percentiles within those bounds. Because the archived clips are trimmed to the selection, duration was obtained directly from the clip length. The pipeline was validated against the 712 annotated signals. It reproduced center frequency and bandwidth with median absolute errors of 21 Hz and 47 Hz, and placed the frequency bounds within a median of 107–147 Hz of the manual bounds, a difference that reflects the subjectivity of selection-box edges and is quantified in Section 3.2. Three of the 20 clips were dominated by broadband noise and contained no measurable call; these were excluded, leaving 17 measured signals and a final dataset of 729.

Acoustic detections were compared with field records of visually observed Bryde’s whales. Detections were assigned to one of three evidence categories: (i) video-confirmed (concurrent video documentation of whales at the surface near the vessel), (ii) visually confirmed by the crew but without video documentation, and (iii) acoustic-only (no concurrent visual confirmation).

### 2.4 Statistical analysis

All statistical analyses were performed in Python 3.13 using NumPy, SciPy, scikit-learn and statsmodels.

Signals recorded within a single session share the same animal, receiver and acoustic conditions, and are therefore not independent observations. Relationships between duration and the frequency parameters were evaluated with linear mixed-effects models including recording session as a random intercept, with both variables standardized (z-scores) so that coefficients are directly comparable. Session means were analysed with Spearman rank correlation as a distribution-free check, and pooled Spearman correlations are reported for comparability with earlier studies. Models were fitted by restricted maximum likelihood. Exact p values are reported throughout, together with rank-biserial effect sizes for the comparison between video-confirmed and unconfirmed signals.

To test whether the measured signals comprise distinguishable call types, the six parameters (duration, MinF, MaxF, center frequency, BW50, BW90) were standardized and subjected to principal component analysis (PCA). Cluster structure was then examined with k-means clustering (k = 2–8; 25 random initializations) and with Ward hierarchical clustering, using silhouette width to assess separation. Cluster stability was quantified with 200 bootstrap resamples, comparing each resampled solution with the full-data solution using the adjusted Rand index (ARI). The biological meaningfulness of the retained solution was evaluated by (i) inspecting cluster centroids in the original parameter units, (ii) testing whether the clusters were present across all recording sessions, and (iii) 5-fold cross-validated linear discriminant analysis (LDA) to quantify separability. The retained three-cluster solution was fitted with 50 random initializations and a fixed random seed, and the clusters were ordered by mean minimum frequency to define Type A, Type B and Type C. Every signal was assigned to this solution, and the resulting classification is provided as supplementary data (Table S1).

The published Bryde’s whale repertoire is defined largely by signal structure, not by spectral parameters alone, so the pulse structure of every signal was also quantified. The band-limited (350–1300 Hz) energy envelope was computed with a 0.89 ms time step, high-pass filtered with a 31-frame median filter, and autocorrelated; the dominant periodicity between 12 and 120 pulses s⁻¹ was taken as the pulse repetition rate, and the corresponding autocorrelation coefficient was retained as a measure of how strongly periodic the envelope was. Frequency-modulation slope was computed from a median-filtered frequency track and is reported as a descriptive measure of signal shape.

Call rate was calculated as the number of signals divided by the time elapsed between the first and last detection in each session, and is therefore a within-session rate.

Noise levels were first assessed by calculating root-mean-square sound pressure levels (SPL_rms_) over the 20 Hz–20 kHz analysis band during periods with and without speedboat passage, using the following formula:

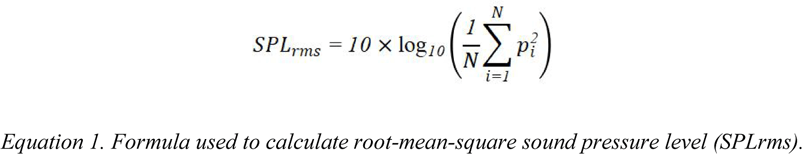

To resolve the spectral distribution of the noise, one-third-octave band levels were computed from five recordings made from the stationary sailboat, totalling 15.7 min and containing nine speedboat passages. Each 1-s window was classified as quiet when its level lay within 1.5 dB of the recording median, and as passage when it lay within 3 dB of the maximum level reached in that recording. This gave 176 quiet and 38 passage windows. Welch power spectral densities were computed for those windows (32,768-sample Hanning window, 50% overlap, 17.6-Hz resolution at the 576-kHz sampling rate) and summed within one-third-octave bands with base-10 centres from 25 Hz to 20 kHz (IEC 61260-1). Levels were referenced to the quiet-condition SPL_rms_ of 119.16 dB re 1 µPa, so that the band analysis and the SPL comparison share one calibration. Communication-range implications were estimated from the call-band level difference assuming transmission loss of the form 10·n·log_10_(R), with n reported for cylindrical (n = 1.5) and spherical (n = 2) spreading.

## 3 Results

### 3.1 Survey effort and occurrence of Bryde’s whale vocalizations

Sixteen surveys were conducted between 11 and 31 January 2024 (21-day survey period). Analysis of the retrieved acoustic data identified signals attributed to Bryde’s whales on eight days: 13, 15, 16, 17, 19, 20 and 21 January, plus a single detection on the night of 14 January (Figure 2; Figure S1). Table 1 lists the ten recording sessions in which signals were measured, with the number of signals, the analysed span, the within-session call rate and the associated evidence category.

**Figure 2.**
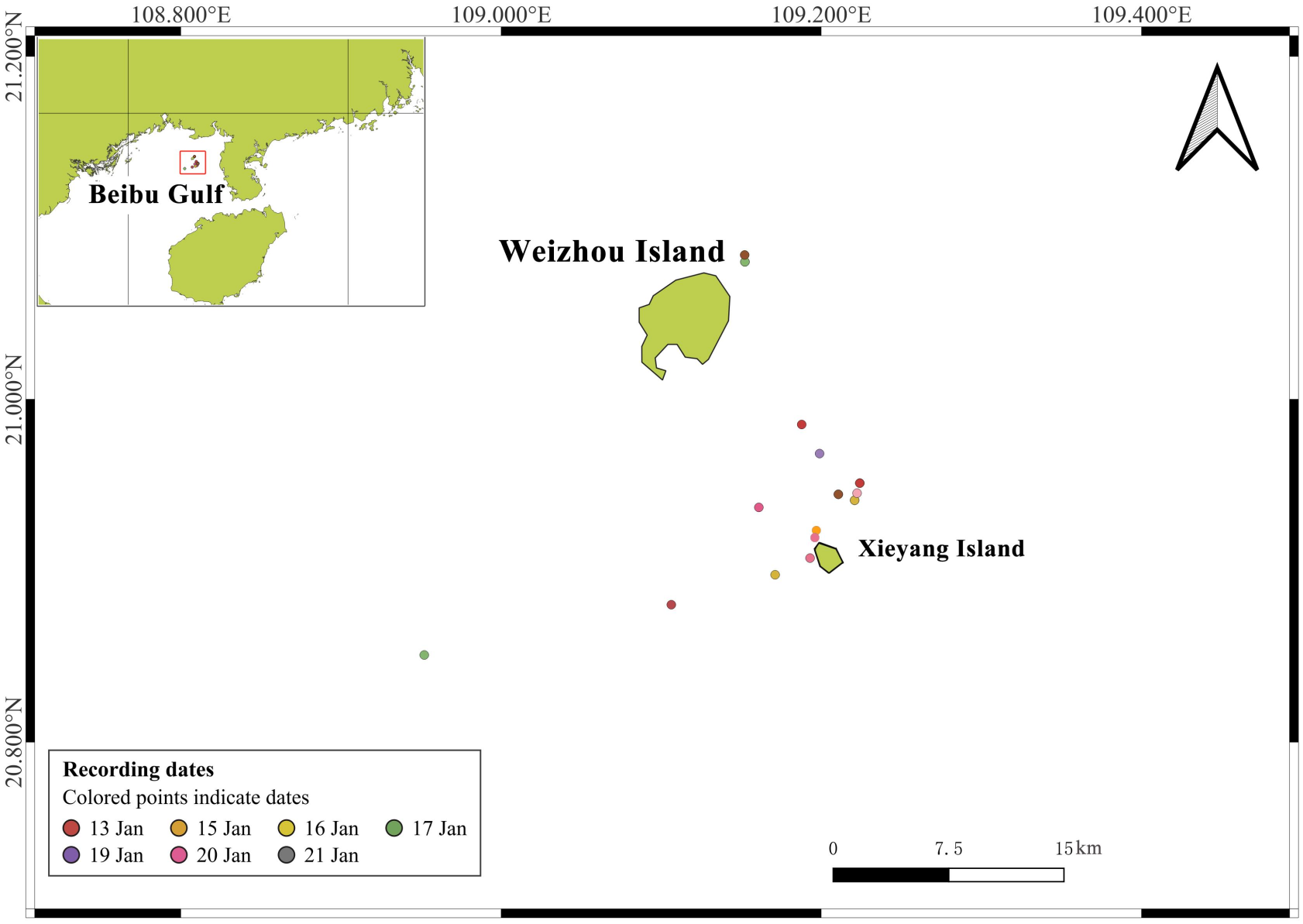
Locations where signals attributed to Bryde’s whales were recorded or observed during the study. Coloured points indicate recording dates. Survey tracks are shown in Figure S1.

Detections were assigned to the three evidence categories defined in Section 2.3. Of the 729 measured signals, 176 (24.1%) were video-confirmed, 343 (47.1%) were recorded during periods when crew members reported visual sightings of Bryde’s whales but no video documentation was available, and 210 (28.8%) were not accompanied by either video documentation or crew sighting reports.

### 3.2 Characteristics of Bryde’s whale vocal signals

We identified 729 low-frequency signals as Bryde’s whale vocalizations according to their spectral structure, recording context and available field sighting records.

Mean duration was 0.73 ± 0.19 s (median = 0.72 s; CV = 25.71%). Minimum frequency averaged 539.56 ± 105.76 Hz (median = 540.76 Hz; CV = 19.60%) and maximum frequency 945.19 ± 110.01 Hz (median = 928.75 Hz; CV = 11.64%), giving a center frequency of 749.42 ± 68.21 Hz (median = 747.07 Hz; CV = 9.10%). The 50% bandwidth was 141.52 ± 49.59 Hz (median = 131.84 Hz; CV = 35.04%), the 90% bandwidth 314.39 ± 103.85 Hz (median = 304.93 Hz; CV = 33.03%), and delta frequency 405.63 ± 147.0 Hz (Table 2). Spectrograms of representative signals attributed to Bryde’s whales and recorded off Weizhou Island are shown in Figure 3.

**Figure 3.**
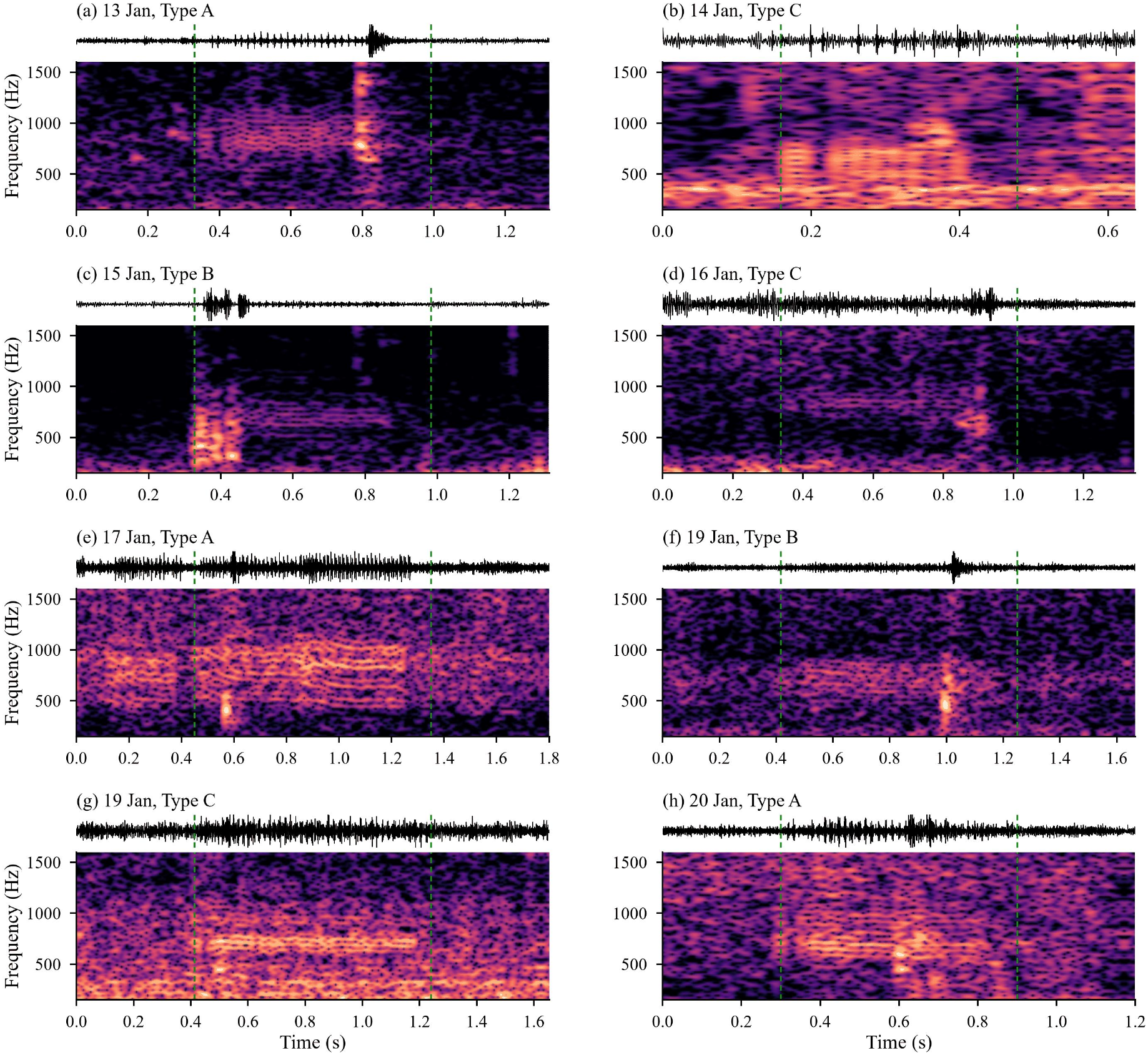
Representative low-frequency signals attributed to Bryde’s whales, recorded off Weizhou Island between 13 and 20 January 2024. Each panel shows a window cut from the continuous 576-kHz recording and centred on one measured signal: waveform (upper strip) and spectrogram (lower). The window is twice the length of the analysed selection, so that each signal is flanked by 50% of its own duration as context; green dashed lines mark the start and end of that selection. One signal is shown for each recording date, with a second signal from 19 January, the session with the highest call density and confirmed visual sightings. Panels are labelled with the recording date and the call type assigned in Section 3.4. Spectrograms were computed with a 32,768-sample Hanning window (17.6-Hz frequency resolution) and 94% overlap (3.56-ms time grid), displayed over 150–1600 Hz and normalised to their own maximum; waveforms are band-pass filtered at 150–1600 Hz for display. These display settings differ from the 65,536-sample window used for parameter measurement (Section 2.3), which was optimised for frequency rather than time resolution.

**Table 2.** Parameters of the low-frequency signals attributed to Bryde’s whales off Weizhou Island (n = 729)

| **Parameter** | **Mean ± SD** | **Median** | **CV (%)** | **Minimum** | **Maximum** |
|---|---|---|---|---|---|
| Duration (s) | 0.73 ± 0.19 | 0.72 | 25.71 | 0.31 | 1.82 |
| Minimum frequency (Hz) | 539.56 ± 105.76 | 540.76 | 19.60 | 173.64 | 831.11 |
| Maximum frequency (Hz) | 945.19 ± 110.02 | 928.75 | 11.64 | 558.86 | 1365.02 |
| Center frequency (Hz) | 749.42 ± 68.21 | 747.07 | 9.10 | 281.25 | 984.38 |
| 50% bandwidth (Hz) | 141.52 ± 49.59 | 131.84 | 35.04 | 19.93 | 421.88 |
| 90% bandwidth (Hz) | 314.39 ± 103.85 | 304.93 | 33.03 | 82.83 | 676.76 |
| Delta frequency (Hz) | 405.63 ± 147.00 | 381.18 | 36.24 | 125.15 | 969.13 |

Restricting the analysis to the 176 video-confirmed signals gave closely comparable bandwidths and slightly lower frequencies: duration 0.67 ± 0.15 s, MinF 504.32 ± 109.94 Hz, MaxF 919.95 ± 92.16 Hz and center frequency 716.86 ± 59.56 Hz, against 0.75 ± 0.20 s, 552.01 ± 102.52 Hz, 956.80 ± 111.71 Hz and 761.60 ± 66.19 Hz for the 553 signals recorded without concurrent video. The confirmed subset therefore reproduces the central tendencies of the pooled dataset: bandwidths were indistinguishable between the two subsets (rank-biserial r = −0.09 and −0.05 for BW50 and BW90), whereas duration and the frequency bounds differed with small-to-moderate effect sizes (r = 0.25–0.44; Mann–Whitney U, p < 0.001). The shift is downward in frequency, as expected for calls recorded at greater range, where high frequencies attenuate first. The 17 signals measured from archived clips showed the same downward shift in their frequency bounds (MinF 466.9 vs 541.3 Hz, MaxF 806.0 vs 948.5 Hz, center frequency 660.5 vs 751.5 Hz; p < 0.001), as expected from the offset of automated bound placement, while their bandwidths were indistinguishable from the annotated population (BW50 p = 0.13; BW90 p = 0.25).

### 3.3 Relationships among signal parameters

Signal duration was negatively correlated with maximum frequency and with center frequency, but not with minimum frequency or with either bandwidth measure (Table 3, Figure 4a). The mixed-effects models, which partition within- and between-session variance, retained both effects (β = −0.129, p < 0.001 for maximum frequency; β = −0.120, p = 0.001 for center frequency). The pooled Spearman coefficients were smaller (ρ = −0.105 and −0.142) because pooling across sessions adds between-session variance that the mixed models remove. Correlations computed on session means (n = 10 sessions) were directionally consistent but did not reach significance, as expected for this number of sessions. The correlation structure among parameters is shown in Figure 4b; as expected, minimum frequency was strongly negatively correlated with BW50 (ρ = −0.52) and BW90 (ρ = −0.66), and BW90 was almost perfectly correlated with delta frequency (ρ = 0.97), confirming that these two variables carry essentially the same information.

**Figure 4.**
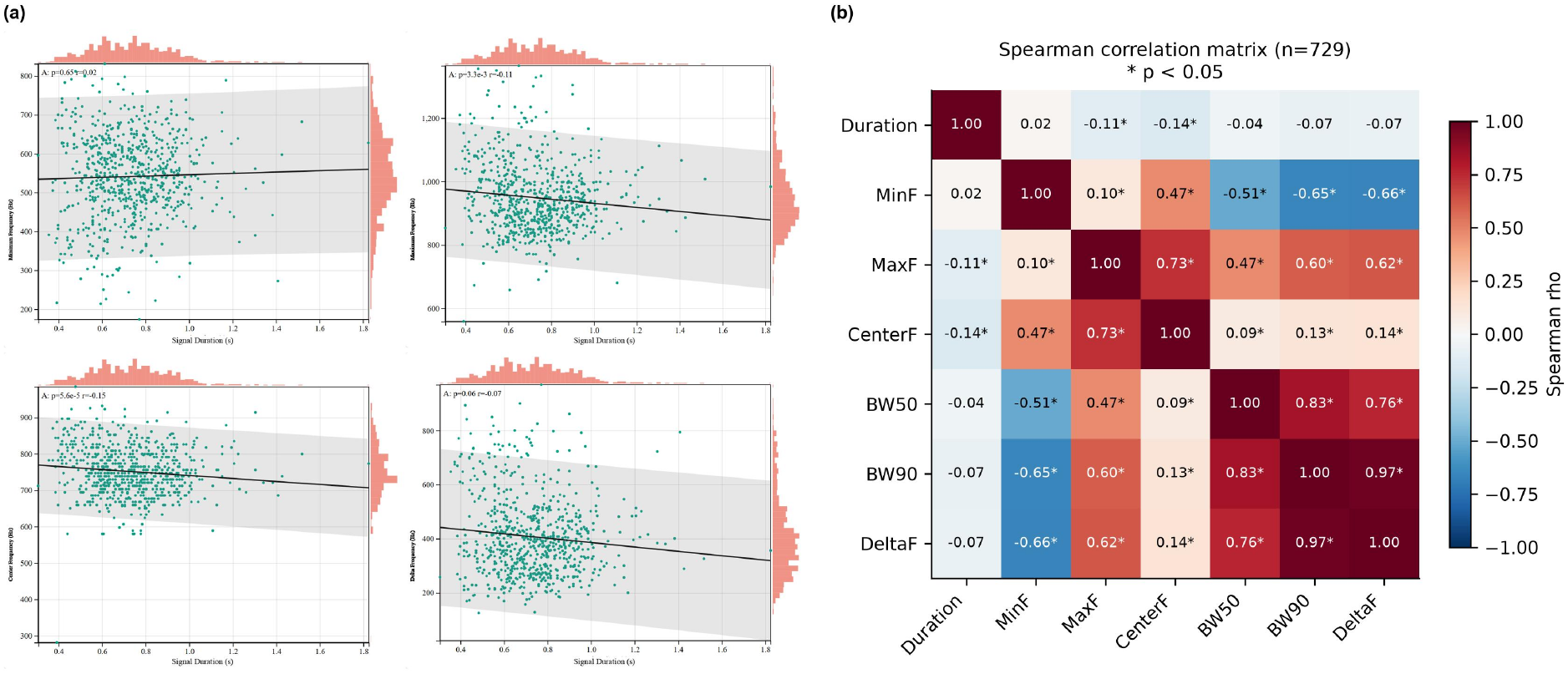
Relationships among signal parameters. (a) Duration against the frequency parameters for the 729 measured signals; (b) Spearman correlation matrix among the seven measured parameters. Asterisks denote p < 0.05.

**Table 3.** Correlations between signal duration and frequency parameters.

| **Parameter** | **Pooled Spearman ρ** | **Pooled p** | **Mixed model β (z)** | **Mixed model p** |
|---|---|---|---|---|
| MinF | +0.017 | 0.642 | −0.044 | 0.239 |
| MaxF | −0.105 | 0.0045 | −0.129 | <0.001 |
| Center frequency | −0.142 | <0.001 | −0.120 | 0.001 |
| BW50 | −0.040 | 0.282 | +0.014 | 0.731 |
| BW90 | −0.071 | 0.056 | −0.048 | 0.219 |

### 3.4 Call-type classification

Every one of the 729 signals had a pulse structure: the band-limited envelope was strongly periodic, with a median pulse repetition rate of 53.6 pulses s⁻¹ (interquartile range 27.4–56.2 pulses s⁻¹; measurable in 669 of 729 signals). The signals therefore belong to the pulsed-call class of the Bryde’s whale repertoire rather than to the tonal or downswept classes that dominate published descriptions of the species (Figure 6).

Within this class, principal component analysis of the six standardized parameters produced three components with eigenvalues greater than one, together accounting for 91.2% of the total variance: PC1 (44.1%) was dominated by bandwidth and upper-frequency loading, PC2 (30.7%) by center frequency and lower-frequency loading, and PC3 (16.4%) by duration.

Silhouette width was positive for k = 2 to 4 and declined smoothly with increasing k (0.303 at k = 2, 0.244 at k = 3), the pattern expected when call parameters vary continuously rather than falling into sharply separated groups. A Gaussian mixture model fitted to the same standardized parameters placed the minimum Bayesian information criterion at k = 3 (8027.9, against 8118.0 at k = 2 and 9037.5 at k = 1; ΔBIC = 1010 for three components over one), with a mean maximum posterior probability of 0.89. Ward hierarchical clustering recovered a congruent topology. The three-group solution was retained because it was stable (mean ARI = 0.87 ± 0.07 across 200 bootstrap resamples), because all three groups recurred across recording sessions (Figure 5c), and because the group centroids differed in parameters with a clear acoustic interpretation.

**Figure 5.**
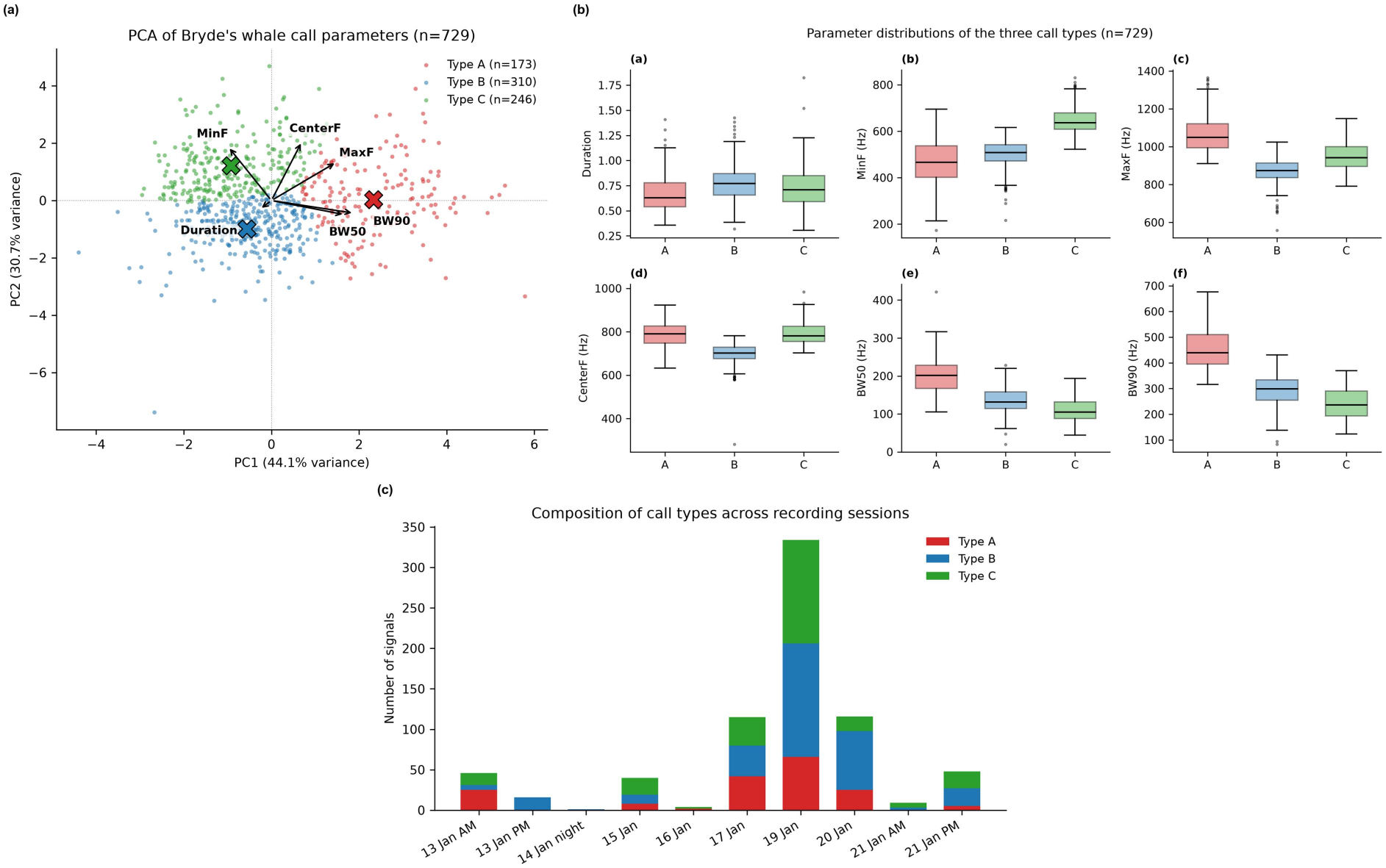
Separation of the three call types. (a) Principal component analysis of the six standardized parameters, with points coloured by call type, crosses marking cluster centroids and arrows showing variable loadings; (b) distributions of the six parameters by call type (i–vi); (c) composition of call types across the ten recording sessions.

A discriminant function fitted to cluster labels recovers those labels whenever the clusters are linearly separable, so the 95.3 ± 3.0% agreement under 5-fold cross-validation measures the internal consistency of the solution rather than providing independent validation. Cluster memberships were exported for all 729 signals (Table S1).

The three call types differed principally in onset frequency and bandwidth (Table 4, Figure 5a,b):

- **Type A (n = 173)** was the broadest-band and highest-reaching type: duration 0.67 s, MinF 462 Hz, MaxF 1069 Hz, center frequency 785 Hz, BW50 200 Hz, BW90 456 Hz.
- **Type B (n = 309)** was the lowest-centred type, with the lowest upper-frequency bound and intermediate bandwidth: duration 0.77 s, MinF 498 Hz, MaxF 872 Hz, center frequency 697 Hz, BW50 135 Hz, BW90 295 Hz.
- **Type C (n = 247)** was characterized by a high onset frequency and the narrowest bandwidth: duration 0.73 s, MinF 646 Hz, MaxF 951 Hz, center frequency 791 Hz, BW50 109 Hz, BW90 239 Hz.

**Table 4.** Parameters of the three call types.

| **Call type** | **n** | **Duration (s)** | **MinF (Hz)** | **MaxF (Hz)** | **Center frequency (Hz)** | **BW50 (Hz)** | **BW90 (Hz)** |
|---|---|---|---|---|---|---|---|
| Type A | 173 | 0.67 | 461.6 | 1069.0 | 784.9 | 199.9 | 456.2 |
| Type B | 309 | 0.77 | 498.3 | 871.5 | 697.0 | 134.6 | 294.8 |
| Type C | 247 | 0.73 | 646.4 | 951.0 | 790.6 | 109.3 | 239.4 |

All three types share the same pulsed structure and differ mainly in where the energy sits within the 400–1100 Hz band (Figure 6). They were present in most recording sessions, although their relative proportions varied (Figure 5c); the 16 measured signals from the afternoon of 13 January were exceptional in that all were assigned to Type B.

**Figure 6.**
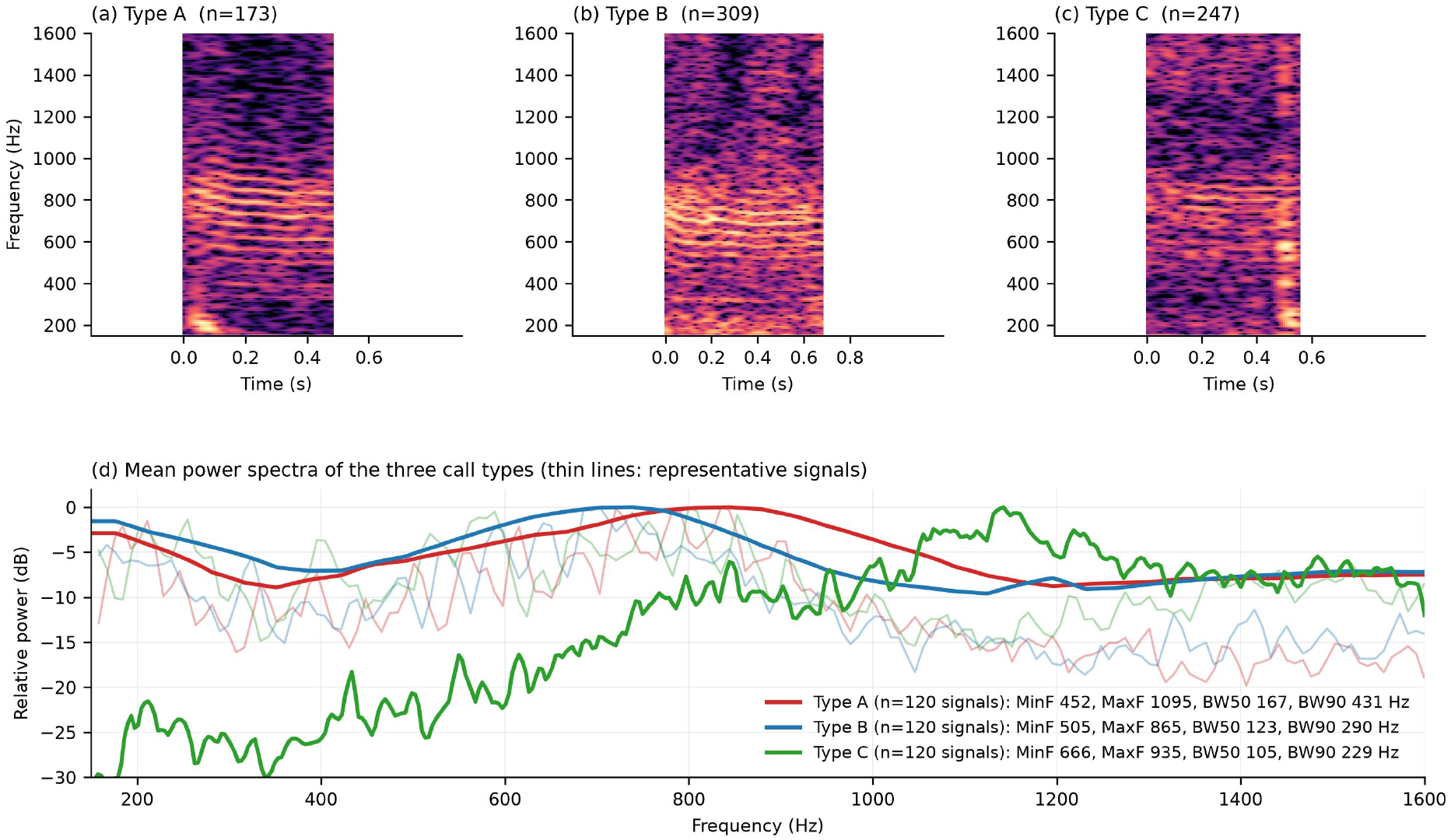
Representative spectrograms (a–c) and type-averaged power spectra (d) of the three call types. Spectrograms were computed with a 65,536-sample Hanning window (8.79-Hz frequency resolution) and 94% overlap, a slightly finer time grid than the 90% overlap used for measurement. The three types differ mainly in where the energy sits within the 400–1100 Hz band; their common pulse repetition rate (20–56 pulses per second) was quantified separately from the band-limited envelope and is not resolved at this frequency resolution.

#### Relation to previously described call types

Table 5 compares the three types with the call classes reported for Bryde’s whales worldwide. The frequency band occupied by the Weizhou Island signals (MinF ≈ 540 Hz, MaxF ≈ 945 Hz) lies at the upper end of the published range for the species, matched only by the growl-like pulsed moan and short pulse sequences described from a captive juvenile and free-ranging adult–calf pairs in the Gulf of Mexico (200–900 Hz, with most energy between 400 and 610 Hz; Edds et al., 1993) and by two of the five call types reported from south-eastern Brazil (Figueiredo and Simão, 2014). In structure, the signals correspond to the discrete-pulse-series class, the most abundant call type in the Brazilian recordings, which was detected while a mother–calf pair was under observation (Figueiredo and Simão, 2014). Within that class, Type A, with its broad bandwidth and high frequency ceiling, most closely resembles the growl-like pulsed moan of Edds et al. (1993), whereas Types B and C, narrower and lower in onset frequency, resemble the short pulse sequences concentrated near 400–610 Hz. The tonal and downswept calls reported from the same region by Wang et al. (2022) (MinF 39–157 Hz, MaxF 105–268 Hz) were not represented in our dataset, and the low-frequency calls reported from the eastern tropical Pacific, the Gulf of Mexico and the South Pacific (Oleson et al., 2003; Heimlich et al., 2005; Širović et al., 2014; Brodie and Dunn, 2015) lie an order of magnitude below our measured frequencies.

**Table 5.** Comparison of the Weizhou Island call types with previously published Bryde’s whale call classes.

| Study | Region | Call class | MinF (Hz) | MaxF (Hz) | Duration (s) | Structure |
|---|---|---|---|---|---|---|
| Cummings et al. (1986) | Gulf of California | short tonal | — | 124 | 0.42 | tonal |
| Edds et al. (1993) | Gulf of Mexico / California | growl-like pulsed moan | 200 | 900 | — | **pulsed** |
| Edds et al. (1993) | Gulf of Mexico / California | short pulse sequence | 400 | 610 | 0.02–0.04 | **pulsed** |
| Oleson et al. (2003) | eastern tropical Pacific | tonal with harmonics | 21–76 | 26–208 | — | tonal |
| Heimlich et al. (2005) | eastern tropical Pacific | tonal phrases | 15–31 | 37–77 | 1.0–2.3 | tonal |
| Širović et al. (2014) | Gulf of Mexico | downswept | 78–85 | 110–143 | 0.4–0.7 | tonal |
| Figueiredo and Simão (2014) | south-eastern Brazil | **discrete pulse series** | — | <1000 | <7 | **pulsed** |
| Figueiredo and Simão (2014) | south-eastern Brazil | flat tone with harmonics | 8.5 | 19 | 1.41 | tonal |
| Figueiredo and Simão (2014) | south-eastern Brazil | downswept tone | 49.5 | 105.5 | 1.16 | tonal |
| Figueiredo and Simão (2014) | south-eastern Brazil | multi-component | 85.7 | 123.6 | 0.88 | complex |
| Figueiredo and Simão (2014) | south-eastern Brazil | multi-component | 225 ± 50 | 555 ± 119 | 0.78 | complex |
| Figueiredo and Simão (2014) | south-eastern Brazil | multi-component | 416 ± 114 | 671 ± 342 | 1.53 | complex |
| Brodie and Dunn (2015) | south-west Pacific | — | 26.1 | 42.3 | 2.5 | tonal |
| Wang et al. (2022) | Beibu Gulf (same region) | repetitive / diversified calls | 39–157 | 105–268 | 0.26–0.55 | tonal, downswept |
| **This study** | **Beibu Gulf** | **Type A** | **462** | **1069** | **0.67** | **pulsed** |
| **This study** | **Beibu Gulf** | **Type B** | **498** | **872** | **0.77** | **pulsed** |
| **This study** | **Beibu Gulf** | **Type C** | **646** | **951** | **0.73** | **pulsed** |

### 3.5 Within-session call rates

Within-session call rates ranged from 0.11 to 5.60 signals min⁻¹ (Figure 7). The highest rates were recorded on 15 January (5.60 signals min⁻¹, n = 40 over 7.1 min), 13 January afternoon (4.92 signals min⁻¹, n = 16 over 3.3 min) and 20 January (4.62 signals min⁻¹, n = 116 over 25.1 min), all sessions with concurrent video documentation of whales near the vessel. The lowest rates occurred on 21 January morning (0.11 signals min⁻¹ over an 81-min span) and 17 January (0.38 signals min⁻¹ over a 305-min span). Median inter-signal intervals ranged from 7.4 s (20 January) to 43.0 s (21 January).

**Figure 7.**
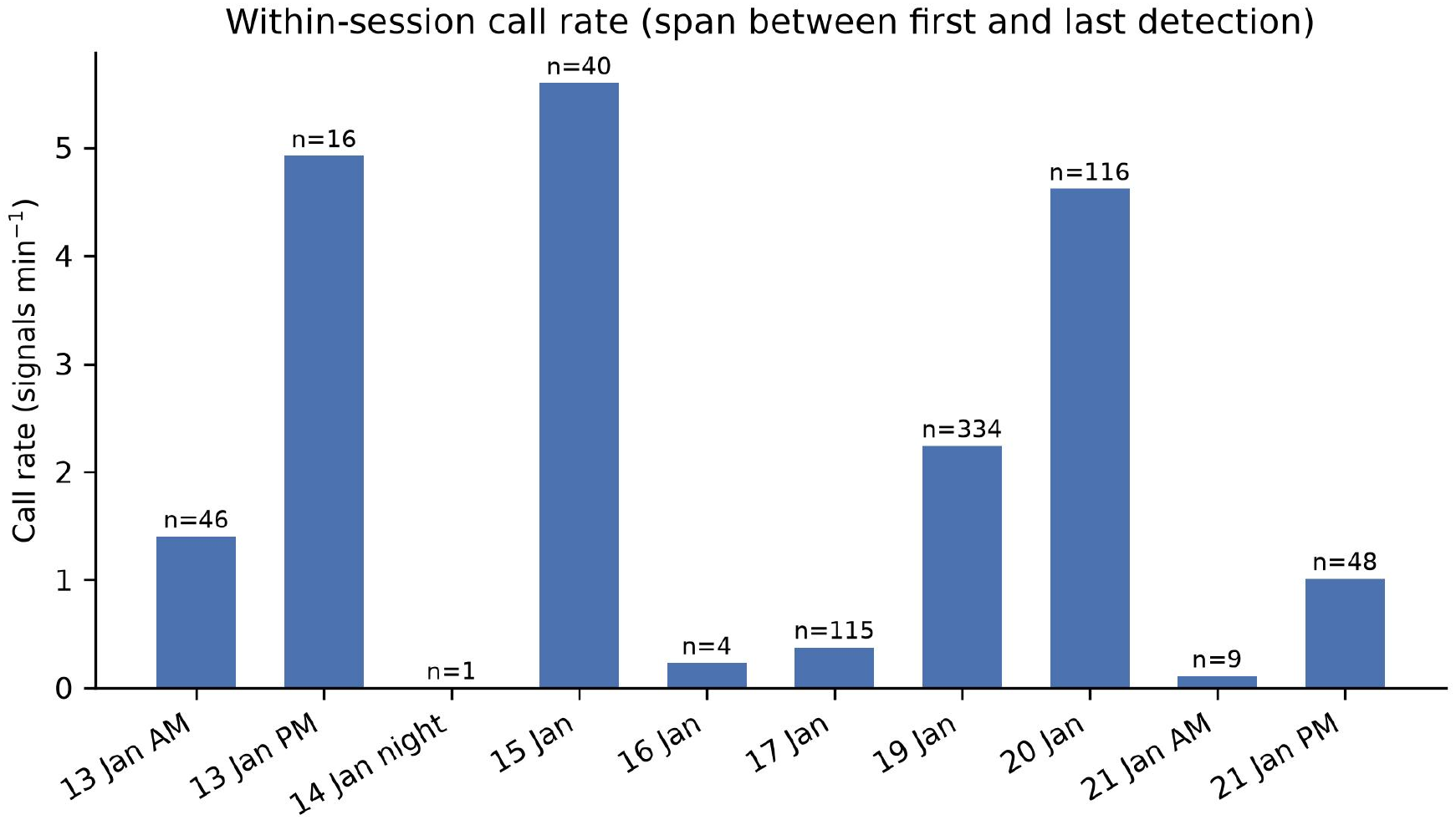
Within-session call rate (signals per minute), calculated over the interval between the first and last measured signal in each session.

### 3.6 Underwater noise associated with speedboats

When the sailboat was stationary, underwater sound was recorded during periods with and without nearby speedboat passage. As shown in Figure 8a, the spectrogram changed markedly during speedboat passage. Sound pressure level was calculated at 1-s resolution over the 20 Hz–20 kHz analysis band, from 15.7 min of recording that contained nine separate passages. During passage the mean level was 130.18 ± 4.63 dB re 1 µPa (median 128.21 dB re 1 µPa; n = 38 windows), against 119.16 ± 2.79 dB re 1 µPa in the absence of speedboats (median 119.07 dB re 1 µPa; n = 176 windows). A Mann–Whitney U test indicated a significant difference between the two conditions (p < 0.0001, rank-biserial r = −0.99; Figure 8b).

**Figure 8.**
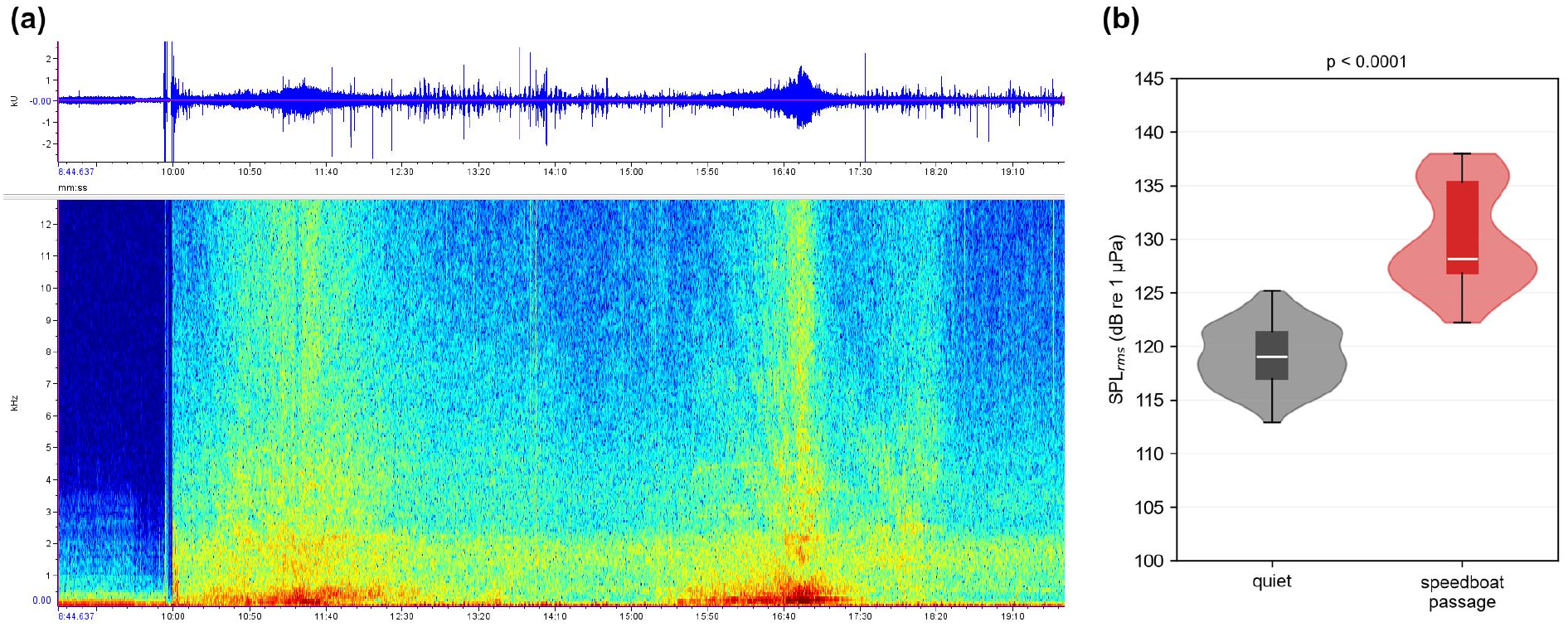
Noise during speedboat passage. (a) Spectrogram of background sound recorded while the sailboat was stationary and a speedboat passed nearby; (b) root-mean-square sound pressure levels with and without speedboat passage. The Mann–Whitney U test indicated a significant difference between conditions (p < 0.0001).

One-third-octave analysis resolved the spectral distribution of this increase (Figure 9). Band-summed over the 20 Hz–20 kHz analysis band the difference was 12.5 dB; this figure is computed from the mean spectra and is therefore slightly larger than the 11.0 dB difference between the mean per-window levels given above. The increase was not uniform across frequency: it was largest below 300 Hz and declined steadily towards the higher bands.

**Figure 9.**
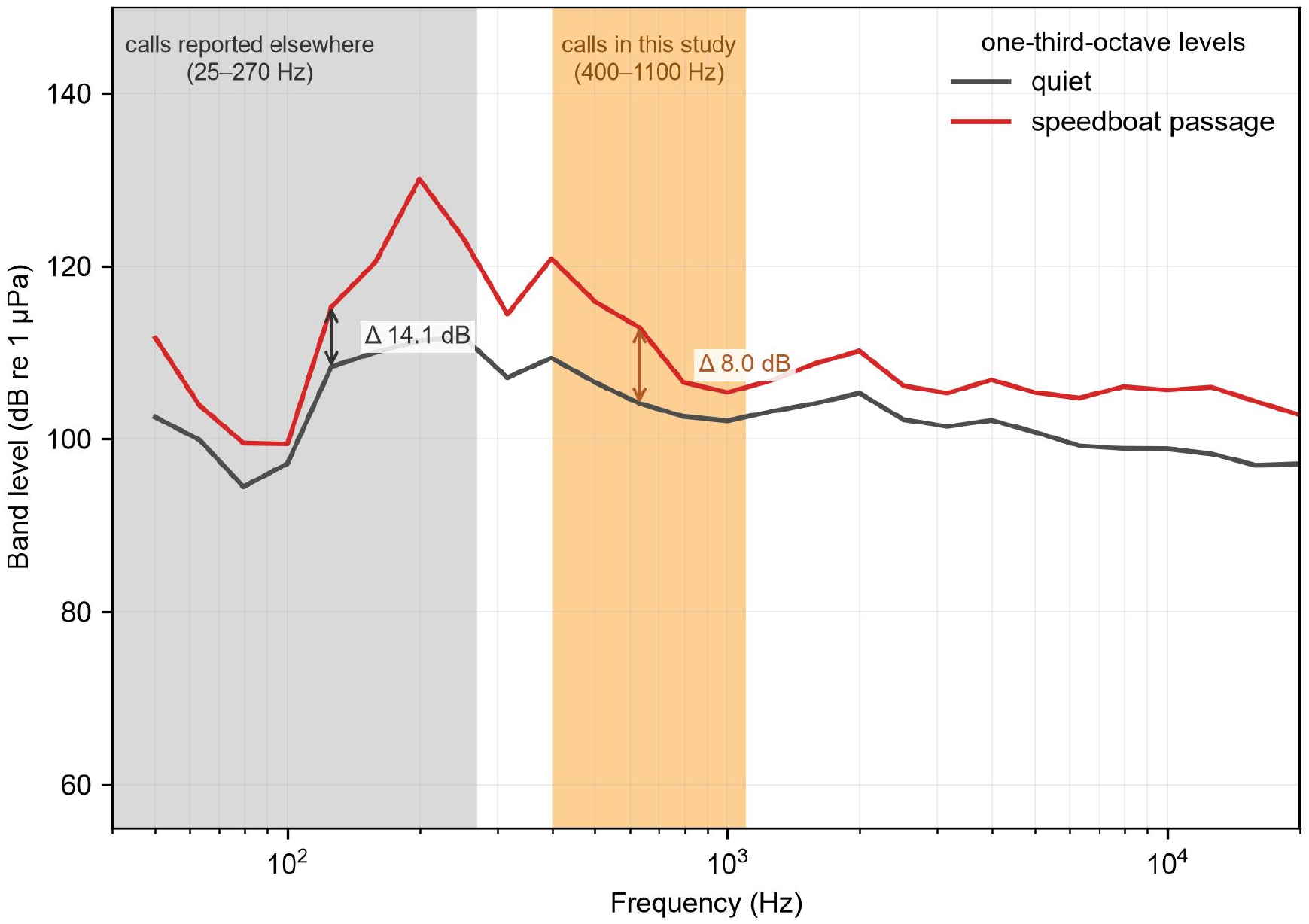
One-third-octave band levels with and without speedboat passage. Shading marks the 25–270 Hz band containing the low-frequency call classes reported for the species elsewhere and the 400–1100 Hz band occupied by the calls recorded here; arrows give the band-summed increase between the two conditions in each band. Levels are calibrated to the quiet-condition SPLrms of 119.16 dB re 1 µPa.

Band levels rose by 14.1 dB in the 25–270 Hz band that contains the low-frequency call classes reported for the species elsewhere, by 9.3 dB in the 400–610 Hz band occupied by the short pulse sequences of Edds et al. (1993), by 8.0 dB in the 400–1100 Hz band occupied by the calls recorded here, and by 4.4 dB between 1.1 and 5 kHz. The noise of a passing speedboat is therefore concentrated at low frequency and overlaps the call band only in part: the calls receive roughly two-thirds of the broadband increase, not the largest share of it.

The consequence for the calls themselves follows from the detection criterion used for annotation. Signals were retained only where the band-limited signal-to-noise ratio exceeded 10 dB, so a call that met that criterion before a passage would, at the same range and source level, fall to about 2 dB above the noise while the boat was passing (10 − 8.0 dB). Masking is therefore partial rather than complete, but it removes most of the margin on which the annotation depended.

If a call of unchanged source level is assumed, an 8.0 dB increase in the noise level within the call band reduces the range over which the call would remain detectable by a factor of about 2.5 under spherical spreading (n = 2) and 3.4 under a cylindrical-to-spherical transition (n = 1.5), that is, to between 29% and 40% of the quiet-condition detection range. The calculation assumes an unchanged source level and a single spreading exponent and does not model absorption or bathymetry.

To place these measurements on a footing comparable with published values, the received levels were converted to equivalent source levels over a range of plausible encounter distances. For an encounter at 25 m the implied broadband source level is 151 dB re 1 µPa at 1 m under cylindrical-to-spherical spreading and 158 dB under spherical spreading, rising to 165 and 176 dB at 200 m. For the most plausible encounters, at 50–200 m, the broadband value is 156–176 dB re 1 µPa at 1 m. Both ranges overlap those reported for small high-speed craft in other coastal waters (Erbe, 2013; Johansson et al., 2024), supporting the view that small planing vessels are governed mainly by engine type, hull form and speed rather than by location, and allowing the levels measured at Weizhou Island to be compared directly with measurements made elsewhere. A directly comparable measurement from the Yangtze River gives a broadband source level of 162.4 dB re 1 µPa at 1 m for a large speedboat travelling at 37 km h⁻¹, against a background of 119.4 dB re 1 µPa (Zhang, 2017). That value lies within the range implied here for an encounter at 25–100 m. Zhang (2017) also reported that speedboat noise energy spans 100 Hz–10 kHz and elevates ambient levels most strongly above 100 Hz, in agreement with the broadband elevation observed in the present recordings. Large merchant vessels radiate at 190–200 dB re 1 µPa at 1 m or more (Erbe et al., 2019), so the speedboats considered here are 20–40 dB quieter than commercial shipping, but they operate in shallow coastal water where the whales occur and where ambient levels are low.

The same arithmetic can be read in the other direction, and doing so constrains the encounter geometry. Combining the published source level of 162.4 dB (Zhang, 2017) with the received level measured here (130.2 dB) gives a transmission loss of 32.2 dB, which corresponds to 41 m under spherical spreading and 139 m under a cylindrical-to-spherical transition; the distance dependence reported by Li et al. (2015) favours the former. The passages recorded here therefore took place within a few tens of metres of the hydrophone, the range at which close-viewing vessels operate and at which masking is strongest.

Table 6 places these estimates alongside direct measurements of small high-speed craft from other coastal and riverine waters, and the agreement is close. High-speed boats in Sanniang Bay, Guangxi, raised ambient levels by 36.9 ± 4.8 dB at 20 m (Li et al., 2015); that figure exceeds the 8.0 dB measured here in the 400–1100 Hz band, which is consistent with their much closer range. The distance dependence reported by Li et al. (2015) is also informative: their boat noise was 8.3 ± 3.8 dB lower at 50 m than at 20 m, which implies a spreading exponent of about 2.1 and therefore supports the spherical-spreading case.

**Table 6.** Noise levels of small high-speed craft reported in previous studies and in the present study.

| Study | Region | Vessel class | Reported metric | Value |
|---|---|---|---|---|
| Zhang (2017) | Yangtze River, China | large speedboat, 37 km h-1 | broadband source level | 162.4 dB re 1 µPa at 1 m |
| Zhang (2017) | Yangtze River, China | small fishing boat, slow | broadband source level | 136.0 dB re 1 µPa at 1 m |
| Li et al. (2015) | Sanniang Bay, Guangxi, China | high-speed boat | received level at 20 m, 6.3 kHz | 149.3 dB re 1 µPa rms |
| Li et al. (2015) | Sanniang Bay, Guangxi, China | high-speed boat | elevation above ambient at 20 m | 36.9 ± 4.8 dB |
| Johansson and Wrede (2026) | Swedish west coast | recreational boat, <10 m | monopole source level vs speed | weak dependence above 7 knots; peak at 5 knots associated with 40–70 Hz tonals |
| **This study** | Beibu Gulf, China | whale-watching speedboats | implied broadband source level | 156–176 dB re 1 µPa at 1 m (50–200 m) |
| **This study** | Beibu Gulf, China | whale-watching speedboats | elevation above ambient | 12.5 dB over 20 Hz–20 kHz; 8.0 dB in 400–1100 Hz |

## 4 Discussion

### 4.1 Bryde’s whale acoustic characteristics and comparison with previous studies

Most published Bryde’s whale calls come from the Gulf of Mexico, the eastern tropical Pacific and south-eastern Brazil (Table 5). Our recordings come from an area where no other baleen whale is known to occur regularly, and none was seen during the survey. The sailboat and manually deployed equipment let us judge recording conditions as they changed and kept engine noise out of the drifting and stationary samples.

The recordings were not made with a localization array, so individual-level attribution was not possible. Bryde’s whales were the only baleen whales observed in the study area, and the signals were repeatedly recorded in temporal and spatial association with sightings, which makes them the most parsimonious source of the recorded vocalizations. The sensitivity analysis in Section 3.2 supports this reading: the video-confirmed subset reproduced the central tendencies of the pooled dataset, with a modest downward shift in frequency consistent with the greater recording range of unconfirmed detections.

The frequencies and durations of the signals recorded here are most similar to calls previously described from captive juvenile Bryde’s whales and free-ranging adult–calf pairs (Edds et al., 1993) and to some call types reported from south-eastern Brazil (Figueiredo and Simão, 2014), which suggests that juveniles and mother–calf pairs contributed to the recordings.

Signal duration was negatively correlated with maximum frequency and center frequency, whereas minimum frequency and bandwidth were unrelated to duration. One plausible explanation is variation among age classes or behavioural states, particularly because ecological studies indicate that Bryde’s whale mother–calf pairs use the waters around Weizhou Island between December and March. Testing this interpretation will require recordings linked to identified juveniles, adults and mother–calf pairs, ideally combining photo-identification with multi-hydrophone localization.

The present dataset also indicates that the vocal repertoire around Weizhou Island is not homogeneous. Three call types were resolved, differing principally in onset frequency, bandwidth and duration, and all three co-occurred across most recording sessions; the cross-validated discriminant accuracy (95.3%) and bootstrap stability (mean ARI 0.87) indicate that the structure is reproducible within this dataset. This bears directly on the comparison with previous work: if the population produces several call classes, the difference between the vocalizations measured here (MinF ≈ 540 Hz, MaxF ≈ 945 Hz) and the lower-frequency calls reported by Wang et al. (2022) (MinF ≈ 40–157 Hz, MaxF ≈ 105–268 Hz) would reflect sampling of different call classes, not a contradiction between datasets. Differences in behavioural context, age composition, recording platform and analysis settings would act in the same direction, and a standardized quantitative call-type framework is the prerequisite for separating them.

The frequency coverage of the dataset is set by the recording configuration. Deployed from a drifting platform, the hydrophone was dominated by flow noise below about 250 Hz, and the analysis was therefore confined to the 400–1100 Hz band in which the calls were detected. The low-frequency tonal and downswept classes reported elsewhere in the species’ repertoire (25–270 Hz; Cummings, 1986; Oleson et al., 2003; Heimlich et al., 2005; Širović et al., 2014; Figueiredo and Simão, 2014; Wang et al., 2022) lie outside this window. Low-frequency calls of this species have been detected with island-mounted seismometers (Xiao et al., 2026), an approach that avoids the flow-noise problem of a drifting platform. Determining whether such calls occur in these waters will require a comparable moored deployment with flow shielding.

The limitations of the dataset follow from its design. The 729 signals come from ten recording sessions, and session was modelled as a random effect throughout. Fifty-seven signals (7.8%) were recorded on 21 January with a second recorder operating at 96 kHz; their bandwidth and duration measurements were consistent with the remainder of the dataset, and they are treated as a single stratum. Seventeen signals were measured from archived clips with the validated automated pipeline, whose bound placement differs from manual annotation by 107–147 Hz; because the classification rests on center frequency, bandwidth and duration, not on the bounds, this offset does not affect the call-type assignment. Night-time recording was restricted to anchorages near Xieyang Island and yielded a single signal on 14 January, so diel patterns remain unresolved.

Close visual presence did not always coincide with acoustic detections: whales sometimes remained within 200 m of the sailboat, or stayed nearby for about 40 min, without any recorded vocalization. Calling therefore occurs in bouts that are not tightly coupled to surface presence, which matters both when acoustic detections are used as a proxy for occurrence and when approach rules are designed around visual cues. Resolving signal direction will require real-time multi-hydrophone arrays.

### 4.2 Potential effects of speedboat noise on Bryde’s whales

Sightseeing speedboats left for whale-watching trips mainly in the afternoon and evening. When the sailboat engine was shut down, we recorded background sound with and without nearby tourist boats. SPL was significantly higher during boat passage (130.18 ± 4.63 dB re 1 µPa; median 128.21 dB re 1 µPa) than during periods without boat passage (119.16 ± 2.79 dB re 1 µPa; median 119.07 dB re 1 µPa; Mann–Whitney U test, p < 0.0001).

The one-third-octave analysis showed that this increase was not uniform across frequency. The largest increments occurred below 300 Hz, where band levels rose by 14.1 dB, and the increase declined with frequency to 8.0 dB in the 400–1100 Hz band and 4.4 dB between 1.1 and 5 kHz. Because the recorded Bryde’s whale signals occupy the 400–1100 Hz band, and because the 90% bandwidth of the three call types spans roughly 240–460 Hz, the increase overlaps the lower part of the band carrying most of the signal energy. This constitutes partial spectral overlap and therefore measurable potential for acoustic masking, which could reduce communication space or require whales to modify calling behaviour, including by increasing call amplitude under the Lombard effect. Direct measurements of mysticete hearing remain confined to a small number of individuals and species (Kleivane et al., 2024), so the consequences for Bryde’s whales are inferred from the overlap between calls and noise rather than from audiograms. Studies of other baleen whales have linked vessel noise with changes in foraging behaviour, migration, stress physiology and vocal behaviour (Blair et al., 2016; Erbe et al., 2018; Helble et al., 2020; Johnston and Painter, 2024; Rolland et al., 2012).

The implied source levels fall within the range reported for small high-speed craft elsewhere (Erbe, 2013; Johansson et al., 2024), so the masking risk documented here is not specific to Weizhou Island: the same classes of vessel operate in whale-watching fleets throughout the region, and the overlap with Bryde’s whale calls would be similar wherever comparable animals occur. The spatial extent of the effect can also be estimated from the measured excess. Relative to the distance at which the measurement was made, broadband noise remained above the pre-passage ambient out to 6–11 times that distance, and noise in the 400–1100 Hz band out to 48–174 times; for an encounter at 50 m this corresponds to roughly 2.4–8.7 km. These are order-of-magnitude estimates: one spreading exponent, no absorption, no bathymetry. Even so, a single passage raises noise well beyond the immediate vicinity of the animal.

Individual speeds were not logged, and the vessels are of a class measured elsewhere at 15–40 km h⁻¹ (Li et al., 2015) and between 3 and 20 knots (Johansson and Wrede, 2026), with individual speedboat passes recorded at 11–37 km h⁻¹ (Zhang, 2017). Because broadband source level changes little above the planing threshold of about 7 knots, with a peak near 5 knots associated with strong 40–70 Hz engine tonals (Johansson and Wrede, 2026), speed is unlikely to alter the levels reported here materially across most of that range; its main effect would be on the low-frequency part of the spectrum, below the band analysed.

The study did not measure behavioural responses to speedboats, and the spectral analysis drew on continuous recordings made on dates other than the SPL comparison. The masking consequences described above are therefore potential rather than demonstrated effects, and pairing behavioural observations with concurrent calibrated noise measurements remains the obvious next step.

Some Bryde’s whales observed during the survey had visible body scars or injuries, consistent with previous reports from this population (Sun et al., 2024; Wu et al., 2020; Zhang, 2021; Zhang et al., 2023). Although the causes cannot be determined from observation alone, the injuries reinforce the case for systematic management of whale-watching activity. Whale watching supports public engagement and conservation awareness, and it can be regulated so that disturbance, collision risk and acoustic exposure are minimized. For Weizhou Island the priorities are evidence-based approach-distance rules, vessel-speed limits, a cap on the number of boats near whales, operator training, and continued monitoring of acoustic conditions and whale behaviour (Figure 10).

**Figure 10.**
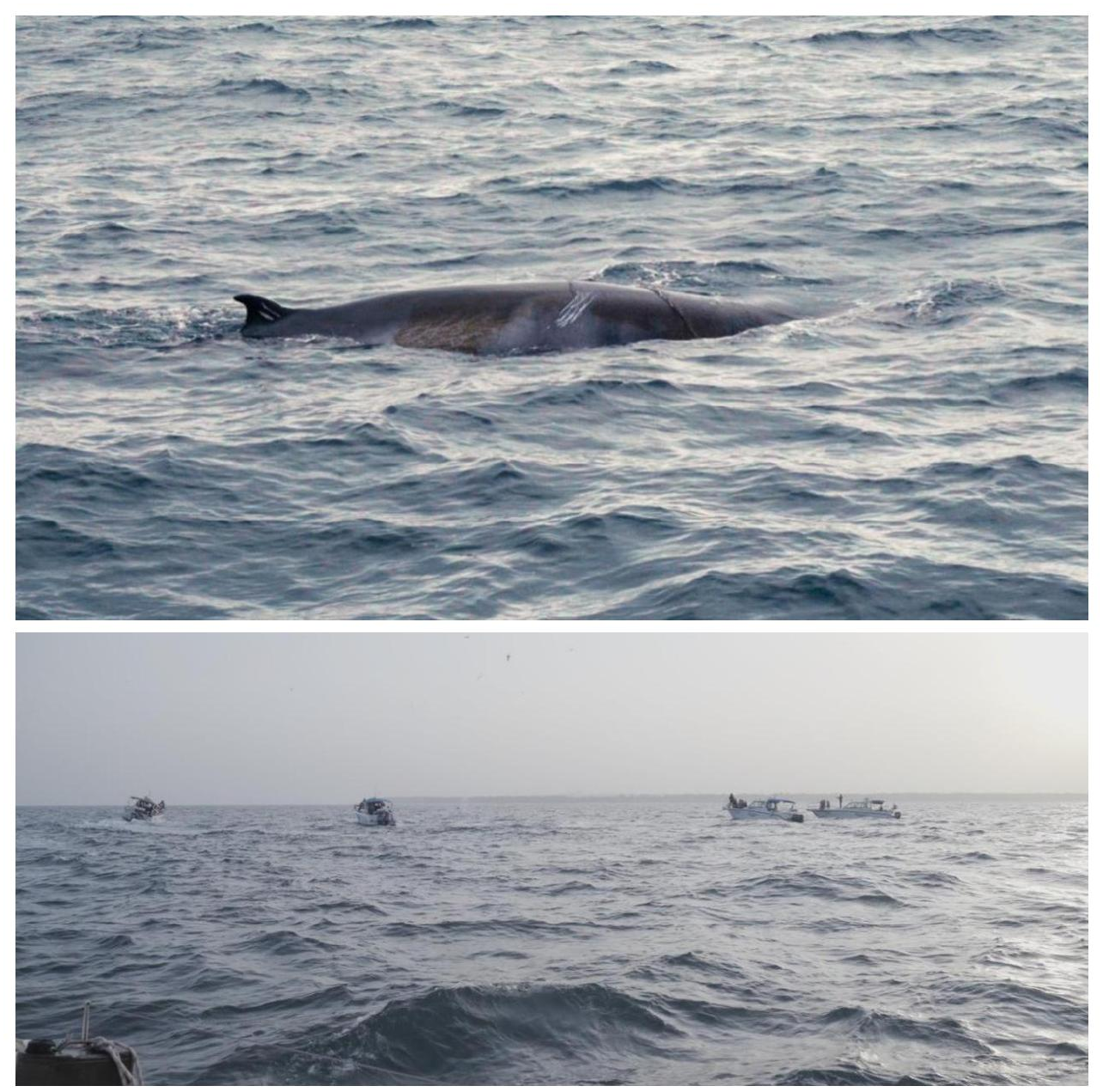
Bryde’s whale with visible scarring photographed during this study (upper panel) and multiple speedboats approaching Bryde’s whales for close viewing (lower panel).

### 4.3 Feasibility of sailboat-based Bryde’s whale surveys

The surveys conducted from 11 to 31 January 2024 show that sailboats can provide a useful platform for Bryde’s whale acoustic surveys around Weizhou Island. The routes overlapped with those used in previous studies (Figure 2; Figure S1) but also allowed access to less frequently surveyed areas. For example, after overnight anchorage near Xieyang Island, whales could be recorded and observed the following morning at locations where their presence had also been documented.

Most previous distribution studies in the region have used speedboat day surveys or questionnaire surveys with fishers, approaches that offer rapid response, relatively low cost and broad coverage (Chen et al., 2019; Sun, 2023; Zhang, 2021). A sailboat cannot respond as quickly, but it can stay on station without an engine running. It also supports day-and-night operations and reaches areas farther from Weizhou Island. In this study, the sailboat reached waters near the oil-platform area approximately 50 km southwest of Weizhou Island, beyond the typical range of small speedboat surveys under many weather conditions. When drifting or operating under sail, the platform also reduced disturbance and improved the likelihood of obtaining usable acoustic recordings near whales.

Sailboat surveys also have logistical limitations. Multi-day offshore operations require a trained crew, including a professional skipper and several crew members capable of sail handling, hydrophone deployment, anchoring and emergency response. Adequate food, freshwater, medical supplies and vessel maintenance support are also required. Future surveys could improve efficiency and safety by using electric windlass systems and electric reefing systems to reduce crew workload during extended field operations.

## 5 Conclusions

A 21-day sailboat survey around Weizhou Island, Guangxi, yielded 729 measured low-frequency signals attributable to Bryde’s whales. All were pulsed, with a repetition rate of 20–56 pulses s⁻¹, and their energy was concentrated in the 400–1100 Hz band, at the upper end of the frequency range reported for the species. The calls most closely resemble the pulsed moans and discrete pulse series previously described from juvenile Bryde’s whales and mother–calf pairs. Cluster analysis resolved three call types that differed in onset frequency, bandwidth and duration and that recurred across most recording sessions; this structure offers a plausible explanation for the discrepancy between the parameters reported here and those previously reported from Weizhou Island, and it expands the acoustic baseline for this coastal population. One-third-octave analysis showed that speedboat passage raised levels by 12.5 dB over the 20 Hz–20 kHz band, with the largest increase below 300 Hz and 8.0 dB within the 400–1100 Hz band occupied by the calls; the implied source levels match measurements of small high-speed craft elsewhere, which places the masking risk beyond this one site. What the calls are used for, and whether vessel traffic changes them, remain open questions. Answering them will need a localization array and behavioural observations tied to identified animals.

## Author contributions

YY: Conceptualization, field investigation, data collection, acoustic analysis, statistical analysis, manuscript preparation. XL: Project coordination, field investigation, data curation, manuscript preparation. ML: Project coordination, field investigation, data curation, manuscript preparation. MC: Project coordination, field investigation, data curation. FF: Conceptualization, project coordination. KW: Conceptualization, project coordination. HD: Project coordination, supervision, project administration, manuscript review and editing.

## Data availability statement

The classification of all 729 signals is provided as Supplementary Material (Table S1). The compiled measurement dataset and the analysis code supporting the findings of this study are available from the corresponding author on reasonable request, as are the raw acoustic recordings.

## Ethics statement

This study relied on passive acoustic monitoring and non-invasive visual observation of free-ranging whales. No animals were handled and no experimental manipulation was conducted. Field procedures followed national guidelines for the observation of free-ranging cetaceans.

## Acknowledgements

The authors thank the field team, vessel crew, local collaborators and supporting institutions for assistance during surveys around Weizhou Island. They are: Shikan Lin, Yaoyao Zhang, Lun Zhang, Tianke Bai, Guang Yang, Peiyi Li, Zhe Zhao, Zhuoxian Luo, Kito, Tao Ju and Lixin Wu. The authors also thank 52Hz Sound Studio and the Institute of Hydrobiology, Chinese Academy of Sciences, for their considerable support of this project.

