## Supplementary figure S1 for "Acoustic Characteristics of Low-Frequency Vocalizations of Bryde’s Whales (*Balaenoptera edeni edeni*) in the Weizhou Island Waters of the Beibu Gulf, China"

*Figure S1. Sailboat survey tracks during the study period (11-31 January 2024). The tracks overlap those used in earlier distribution studies and extend to waters near the oil and gas platforms southwest of Weizhou Island.*

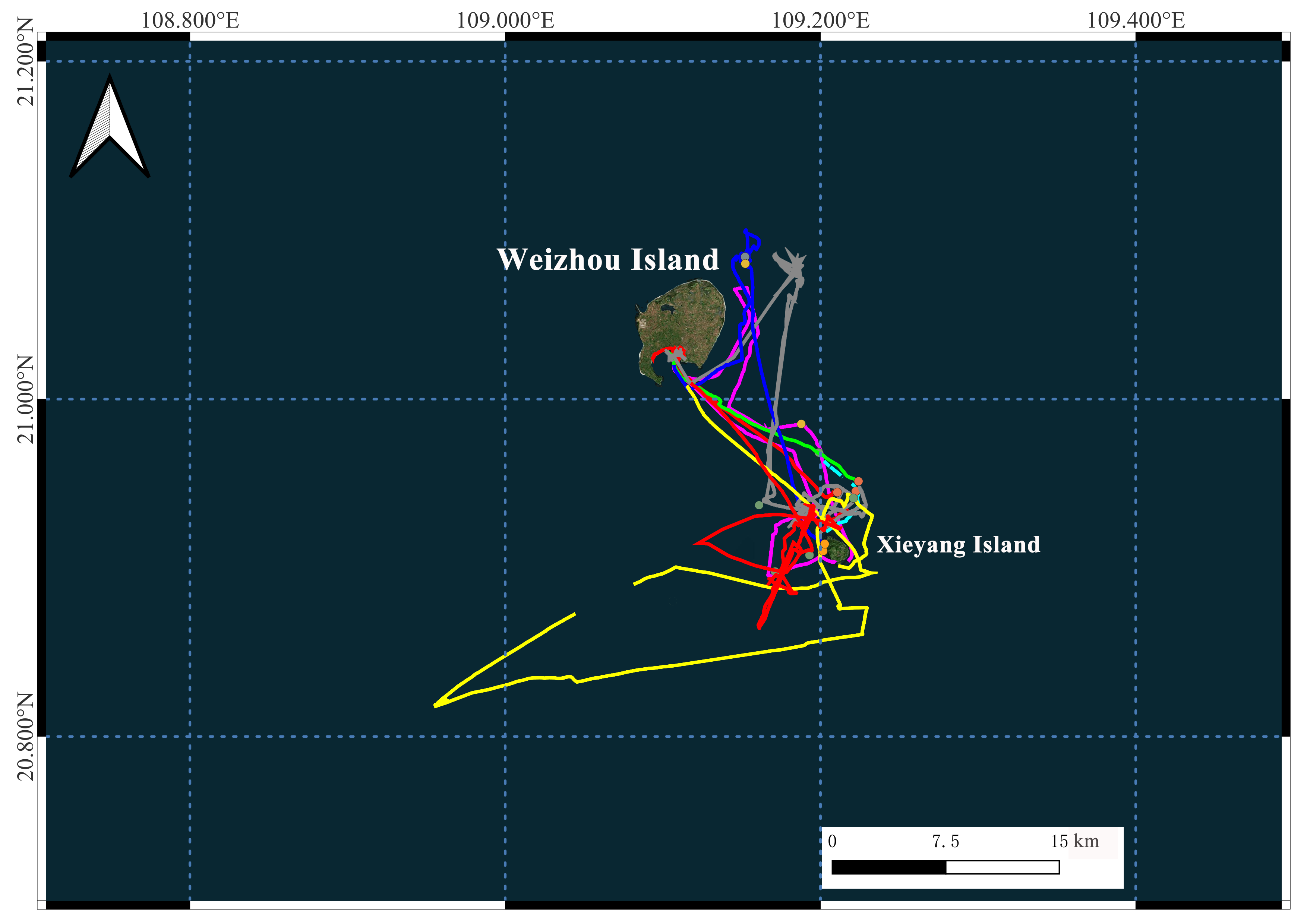

**Note on data availability**

The measurement dataset and the analysis code are available from the corresponding author on reasonable request, as are the raw acoustic recordings.
